# Illumina MiSeq, MiSeq i100, and Element AVITI yield concordant ecological patterns for universal 16S and 18S rRNA gene amplicon sequencing

**DOI:** 10.64898/2026.11.14.766689

**Authors:** Elisa D’Agostino, David M. Needham

## Abstract

Molecular microbiome analyses require high-resolution, comparable measurements, often obtained by sequencing marker genes such as the rRNA gene. Recently, new platforms and chemistries have promised to improve performance at lower cost relative to the commonly used Illumina MiSeq 2×300, remains largely untested especially for rRNA gene amplicon studies. Here, we compared MiSeq 2×300, MiSeq 2×500, and AVITI 2×300 by sequencing mock communities and field samples via a ‘universal’ (16S + 18S) rRNA gene amplicons (V4-V5 region) approach. All platforms recovered similar community patterns (Mantel r ≥ 0.93), with no major differences in recovered lineages. MiSeq 2×500 showed ∼4.5-fold higher index hopping (5.9 ± 3.8% vs. 1.3 ± 1.5%). For 18S, MiSeq 2×500, despite longer reads overlapping the 18S V4-V5 region, only marginally improved taxonomic classification (+0.7% on average) and ASV resolution (+1.3%). Meanwhile, all platforms similarly biased against 18S due to its greater length, which we demonstrate can be offset by a gel-based size selection, with no apparent taxonomic bias. Overall the platforms yield ecologically similar results. We highlight platform-specific advantages to help guide decision-making and challenges that can be addressed through dedicated bioinformatic or wet-lab approaches.

## INTRODUCTION

Molecular characterizations of biodiversity are ideally quantitative, precise, and comparable across studies and datasets (Fuhrman et al., 2015; Lima-Mendez et al., 2015; Worden et al., 2015; Rao et al., 2021; Shaffer et al., 2022), to enable insights and meta-analyses across systems and over long-term time series. For over a decade, amplicon sequencing studies have largely depended on one sequencing technology, Illumina MiSeq, with remarkable success. Recently, two new platforms that promise to complement MiSeq have emerged: the MiSeq i100 2×500 (hereafter, “MiSeq 2×500”), which allows for longer reads and more throughput, and the Element Biosciences, AVITI (hereafter, AVITI 2×300), which offers higher quality (Arslan et al., 2024; Liu et al., 2024) and throughput.

Here, we compared the performance of these sequencing platforms using a ‘universal’ rRNA gene sequencing approach. This approach utilizes the 515F/926R primer pair targeting the V4-V5 hypervariable regions (Parada et al., 2016) and enables sequencing of prokaryotes and eukaryotes in a single ‘universal’ assay (Needham and Fuhrman, 2016; Parada et al., 2016) that performs as well or better for each individual domain as domain-specific primers (Stoeck et al., 2010; Klindworth et al., 2012; Balzano and Abs, 2015; Mahé et al., 2015; McNichol et al., 2021; Yeh et al., 2021). For this primer set, notably, 18S rRNA gene amplicon (hereafter, “18S”) assessment has been limited by the low relative proportions of eukaryotes vs. prokaryotes in many environments (Whitman et al., 1998; Bar-On et al., 2018) and by the 18S being ∼200 bp longer than the 16S rRNA gene amplicon (hereafter, “16S”), which biases against 18S during PCR (Yeh et al., 2021; Shaffer et al., 2025) and Illumina sequencing (Kittelmann et al., 2013; Gohl et al., 2019; Yeh et al., 2021). This length also complicates bioinformatics workflows because the two reads do not overlap across the 18S V4-V5 region when assessed by 2×300 sequencing (Needham and Fuhrman, 2016; Ramakodi, 2024; Priest et al., 2025). The MiSeq 2×500 can help address this as it offers overlapping sequences for the 18S, while AVITI’s lower cost per read relative to the other sequencers may offer advantages to robustly detecting rare taxa, including microbial eukaryotes in microbiome samples dominated by prokaryotes.

For assessment of the different sequencing platforms and chemistries, we used 16 mock communities composed of synthetic templates representing sequences from marine microorganisms from all domains of life, alongside 165 marine and brackish field samples from the North Sea and Elbe River, and 22 no-template controls (blanks). We sequenced all of the libraries on Illumina MiSeq 2×300, MiSeq 2×500, and AVITI 2×300. For AVITI 2×300, in addition to the standard approach, we assessed a gel-excision-based method to increase the amount of 18S relative to 16S in the sample (AVITI 2×300 size-selection); there, the proportion of 18S was raised from 17% to 50%, a 2.9-fold increase. Thus, in total, we sequenced the same libraries across the three platforms, plus a fourth run for the AVITI 2×300 size-selection (Supplementary Figure 1).

## EXPERIMENTAL PROCEDURES

### Sample collection

Seawater and brackish water samples were collected across cruises from the German Bight and the Elbe River in 2024 and 2025. Water was filtered on 0.2 µm filters (0.2 µm Supor™ 200 Membrane Disc Filters) with a volume ranging between 200 and 250 mL, typically from the surface (top 1 m) but also including some bottom depths (Supplementary Table 1). 3 µm size fraction samples were gravity-filtered on 3 µm Nucleopore filters with volumes ranging from 100 to 250 mL. Filters were stored in microtubes pre-filled with mixed glass beads (0.1 mm and 0.5 mm) at -80 °C for subsequent DNA extraction.

For the mock community, five bacterial, five eukaryotic, and two archaeal rRNA gene sequences synthesised from Twist Bioscience were pooled at either equimolar or staggered concentrations (Supplementary Table 2).

### DNA extraction

DNA was extracted from each filter using a modified DNeasy Plant Kit (Qiagen) protocol, as previously described (Ghotbi et al., 2025; Kim et al., 2026). Briefly, lysis buffer from the kit (AP1) was added to each microtube with the filter and beads, followed by three freeze-thaw cycles with liquid nitrogen and a 60 °C water bath. Mechanical lysis was performed via bead-beating using a Laboratory Mixer (MM400, Retsch). The lysate was then transferred to a 96-well plate for purification via an AcroPrep plate. A total of 14 blanks were included and distributed randomly across extraction plates. Additionally, spike-in controls (ZymoBIOMICS Spike-in Control I, High Microbial Load, D6320) were added to each sample prior to DNA extraction.

DNA was eluted in Tris-EDTA and stored at -80 °C until amplicon library prep.

### PCR amplification

For amplicon libraries, 1 ng and 0.5 ng were input into the PCR for the 0.2 and 3 μm size fractions, respectively. In addition to the 14 extraction blanks, eight no template PCR controls were also included. The universal 515F/926R primer set targeting the V4-V5 hypervariable region (Parada et al. 2016) was used. The V4-V5 primers were synthesised to include Illumina P5 adapter sequence, a unique 8-bp index, sequencing primer, and the template-specific primer which is added via a single PCR reaction of 30 cycles, as previously described (Ghotbi et al., 2025; Kim et al., 2026). Mock communities (even and staggered) were added to the PCR plates. Two mock community samples (29Oct24 and 23Oct25-4, even and staggered) were excluded from subsequent analyses due to poor recovery of the expected inputs, which were consistent with wet-lab handling error (Supplementary Table 1). After PCR, amplification was assessed via gel electrophoresis. The amplified products were purified via a 0.8× Ampure XP magnetic bead clean-up and quantified with Quant-iT™ PicoGreen™ dsDNA Assay, and then pooled in approximately equimolar concentrations. An aliquot of this pool was directly sequenced with MiSeq 2×300 and MiSeq 2×500 at the Competence Centre for Genomic Analysis (Kiel, Germany). For AVITI 2×300, out of the equimolar pool, 150 μL were taken and purified via an additional 0.8× bead clean-up, with a final elution of 75 μL, which underwent an “End-Polishing” PCR prior to sequencing, see below.

### Size-selection

For size-selected libraries via gel extraction, on the remaining volume of the equimolar pool, a 1x bead clean-up with final elution of 25 μL was performed. This DNA was then distributed into two lanes of a 2% agarose gel in a 1× TAE buffer at 75 volts for one hour. The gel was inspected for clear separation of the 16S and 18S bands and then excised with a sterile scalpel, purified from the gel via PureLink™ Quick Gel Extraction kit (Invitrogen) and finally eluted in 50 µL. Complete separation of the two bands from the excised products was confirmed via BioAnalyzer DNA12000 and the average size of the 16S and 18S bands was determined. Quantitation of the libraries was also checked with Qubit, and molarity calculated considering the average length of the product.

### AVITI End-Polishing

Due to guidance that AVITI sequencing is sensitive to incomplete or truncated adapters, the gel extracted 16S and 18S pools as well as the original equimolar pool were “End-Polished” with PCR with the P5 and P7 primers. For each of the two pools, 5 ng were added to separate PCR reactions and amplified as follows: initial denaturation at 98°C for 90 seconds; 5 cycles of denaturation at 98°C for 30 seconds, annealing at 60°C for 30 seconds, and extension at 72°C for 30 seconds; final extension at 72°C for 1 minute. Fragment analysis for AVITI was determined with Fragment Analyzer, and BioAnalyzer DNA12000 was run on 0.8× Ampure bead-cleaned individual PCR reactions of staggered and mock communities. AVITI sequencing was carried out at Lexogen (Vienna, Austria).

### Bioinformatics

For 2×300 sequencing runs, V4-V5 amplicons, primers 515F and 926R were trimmed from raw reads using Cutadapt v4.4 (Martin, 2011) (-g GTGYCAGCMGCCGCGGTAA -G CCGYCAATTYMTTTRAGTTT) with maximum 0.2 errors (-e 0.2) and no indels (--no-indels) and discarding untrimmed sequences (--discard-untrimmed). 16S and 18S amplicons were split with bbsplit.sh (Bushnell, 2014) by comparison to SILVA 132 (Quast et al., 2013) and PR2 (Guillou et al., 2013). 18S reads were pasted together and then both 16S and 18S denoised (separately) with the DADA2 plug-in for QIIME2 (Callahan et al., 2016). For 2×500 sequencing, raw reads were again trimmed with Cutadapt, but allowing for trimming on the 3’ end for the opposing primer to prevent read-through (-a “GTGYCAGCMGCCGCGGTAA…AAACTYAAAKRAATTGRCGG”, -A “CCGYCAATTYMTTTRAGTTT…TTACCGCGGCKGCTGRCAC”), allowing 0.1 errors, a minimum length of 150 bp (-m 150), and trimming reads to maximum length of 450 bp). Then, 16S and 18S sequences were processed together by denoising via the DADA2 plug-in in QIIME2 allowing maximum 2 expected error on forward and reverse reads (--p-max-ee-f 2, --p-max-ee-r 2) with 0 truncation (--p-trunc-len-r 0, --p-trunc-len-f 0), and discarding untrimmed sequences (--discard-untrimmed). Index hopping was estimated by counting the number of reads in blanks relative to samples.

Taxonomic classification of ASVs was performed using the *sklearn* classifier within QIIME2 (Pedregosa et al., 2011) against the SILVA 138.1 database to partition sequences into Prokaryotes (16S), Chloroplasts (16S), and Eukaryotes (18S). Subsequently, *sklearn* classifications using Greengenes2 (McDonald et al., 2024) were applied to Prokaryotes, PhytoRef (Decelle et al., 2015) to Chloroplasts, and PR2 (Guillou et al., 2013) to Eukaryotes.

In order to assess whether 2×500 18S sequencing provides improved taxonomic assignment or higher ASV resolution, we performed a direct comparison with 2×300 data. To enable this comparison, denoised and stitched (pasted) 2×300 reads that formed ASVs were partitioned into their constituent 250 bp and 200 bp segments. These segments were then searched for exact matches against the overlapping, denoised 2×500 18S ASVs, resulting in “perfect F/R matches”. We counted the number of 2×500 ASVs matching each pair of segments to determine whether additional resolution was achieved with 2×500 sequencing. Likewise, we also evaluated the taxonomy assignment of “perfect F/R matches” by comparing the taxonomy assigned to the 2×300 ASVs with that of the corresponding 2×500 amplicons.

All downstream analyses were performed in R (version 4.2.2). Correlation between observed and expected mock communities was calculated by ordinary least squares (OLS) regression (base R::lm) for staggered communities, and Bray-Curtis via vegan::vegdist for both staggered and even. The *vegan* package (v. 2.6-6.1) was also used for Mantel test comparisons, including Bray-Curtis, Jaccard, Hellinger and Aitchison dissimilarities (via vegan::vegdist). *ggplot2* (v. 4.0.2) was used for data visualization.

## RESULTS

We used the mock communities to compare platforms across a known input of taxa distributed in an ‘even’ format (12 taxa, 8.33% each) and ‘staggered’ format (12 taxa, ranging 0.001 to 34.6%). At the community level, within the measured 16S and 18S, across platforms the overall similarity was high, 0.92-0.96 for even samples and 0.94-0.97 for staggered. Compared to the input communities, the measured mock communities were similarly different from the known inputs, with Bray-Curtis similarity to expectation of 0.65– 0.71 (Fig., 1a, even) and 0.80–0.85 (Fig. 1b, staggered), the main deviation being one 18S ASV, *Lingulodinium*, which had a single mismatch to the reverse primer (as noted previously (Yeh et al., 2021)) and was poorly recovered across all platforms (i.e., a difference driven by a primer incompatibility conserved across sequencing platforms). Across all platforms, overall 18S was under-represented relative to input (even: 21.1 ± 2.8% vs 36.4% expected; staggered: 29.4 ± 4.6% vs 40.96%, Fig. 1), consistent also with past results for MiSeq 2×300 (Yeh et al., 2021; McNichol et al., 2025). With the size-selected library, the proportion of 18S was increased by ∼1.8-fold on average (even: 39.7 ± 5.1%; staggered: 51.6 ± 7.5%), about half of the enrichment originally achieved during the size-selection process, indicating a bias against 18S also in the size-selected libraries. Overall, the mock community results indicate that despite differences in the expected inputs due to primer bias or mock community generation, the recovery of mock communities is highly similar across the different platforms.

**Figure 1.**
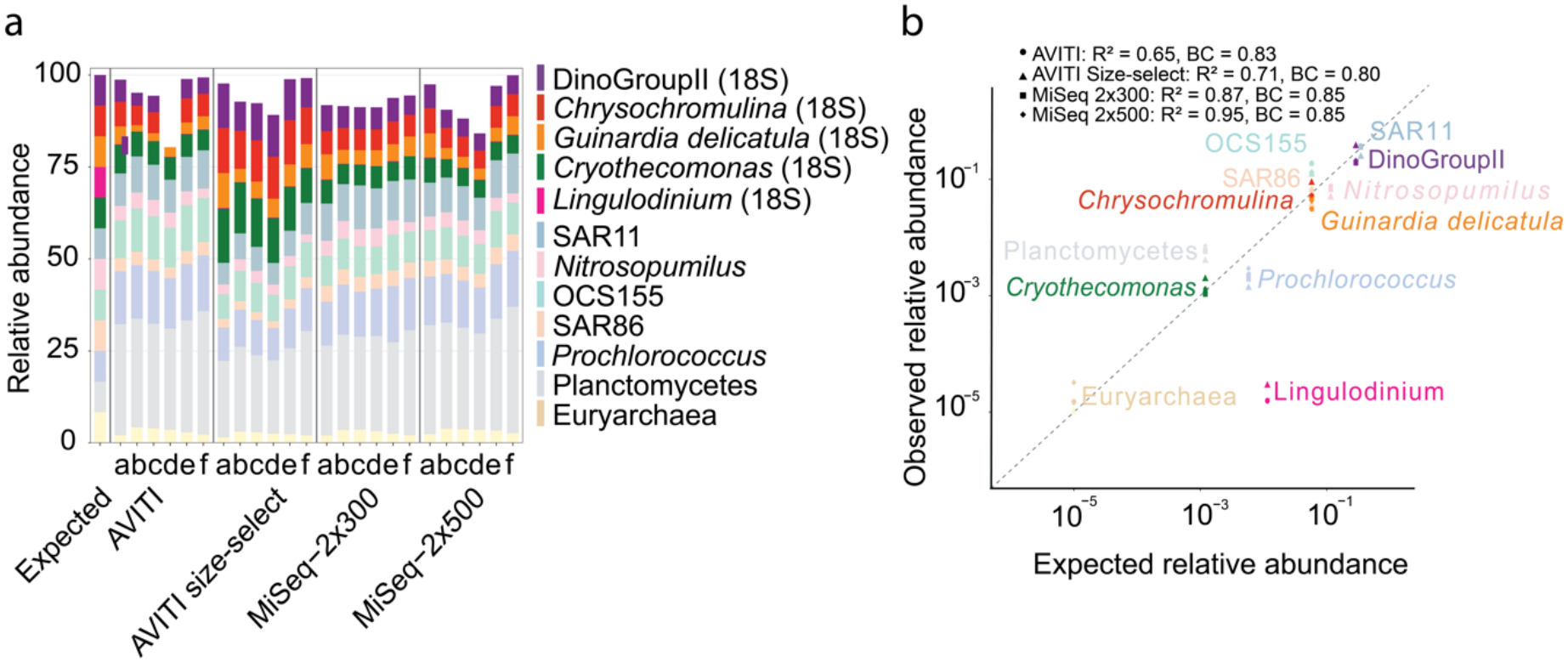
Comparison of results across platforms and chemistries for mock communities. a, Observed vs expected relative abundance of taxa in the even mock community across the four sequencing conditions (n = 6 replicates, as indicated by a-f). The relative abundance values do not equal 100% because of the detection of non-input ASVs in the sequenced mock communities. Mock community similarity calculations included all ASVs detected in the data regardless of whether the ASVs were in the original input, to account for error due to index-hopped sequences and/or sequencing error differences across platforms. b, Observed vs. expected scatter plot for the staggered mock community. Replicates (n=6) are averaged. The dashed line is a 1-to-1 line. Note axes are log-scaled. BC = Bray-Curtis similarity.

For field samples, sequencing results were highly concordant across platforms and chemistries. Overall, the relative proportions of ASVs classified at the Class level for both 16S and 18S were highly correlated across all sequencing platforms (R^2^=0.955-0.992 and R^2^=0.93-0.96, respectively, Supplementary Table 3). All the platforms and approaches showed highly similar ecological patterns as evidenced by strong Mantel correlations at both 18S and 16S (Mantel r ≥ 0.93, p = 0.001 for all pairwise comparisons, Supplementary Table 4). Differences were consistent across different metrics, with Jaccard, Hellinger and Aitchison distances showing similar pairwise patterns (overall Mantel r > 0.84, p = 0.001, Supplementary Table 4), with the lowest correlations tending to be via Hellinger and Aitchison, suggesting that rare taxa are contributing more variability than more abundant taxa. Following up on the impacts of reduced sequencing depth, we compared similarity of communities across abundance categories (dominant, intermediate, rare, very rare; adapted from (Logares et al., 2014)) for all pairwise platform comparisons (Fig. 2a). Similarity remained high across all abundance categories for platform pairs except those with MiSeq 2×300 (Bray-Curtis similarity ≥0.71 and ≥0.89 for very rare 18S and 16S taxa, respectively). Comparisons involving MiSeq 2×300 showed reduced similarity for rare and very rare taxa (as low as 0.33 for 18S). This indicates that the signal obtained from AVITI 2×300 and MiSeq 2×500 higher sequencing depth, indeed improves the recovery of rare taxa for both 16S and 18S. We also noted the lowest Mantel often also corresponded to the MiSeq 2×500. We reasoned that this could be a consequence of increased index hopping in MiSeq 2×500, which is a consequence of its patterned flow cell type (Illumina, 2017; Flörl et al., 2026). We aimed to quantify this by proxy: i.e., counting the number of reads observed in no template controls relative to real samples or mock communities. This metric suggested that index-hopping increased by ∼4.5-fold on MiSeq 2×500 relative to the other platforms (5.9 ± 3.8% vs 1.3 ± 1.5%, Fig 2b), which may help explain some relatively lower Mantel test values for the MiSeq 2×500.

**Figure 2.**
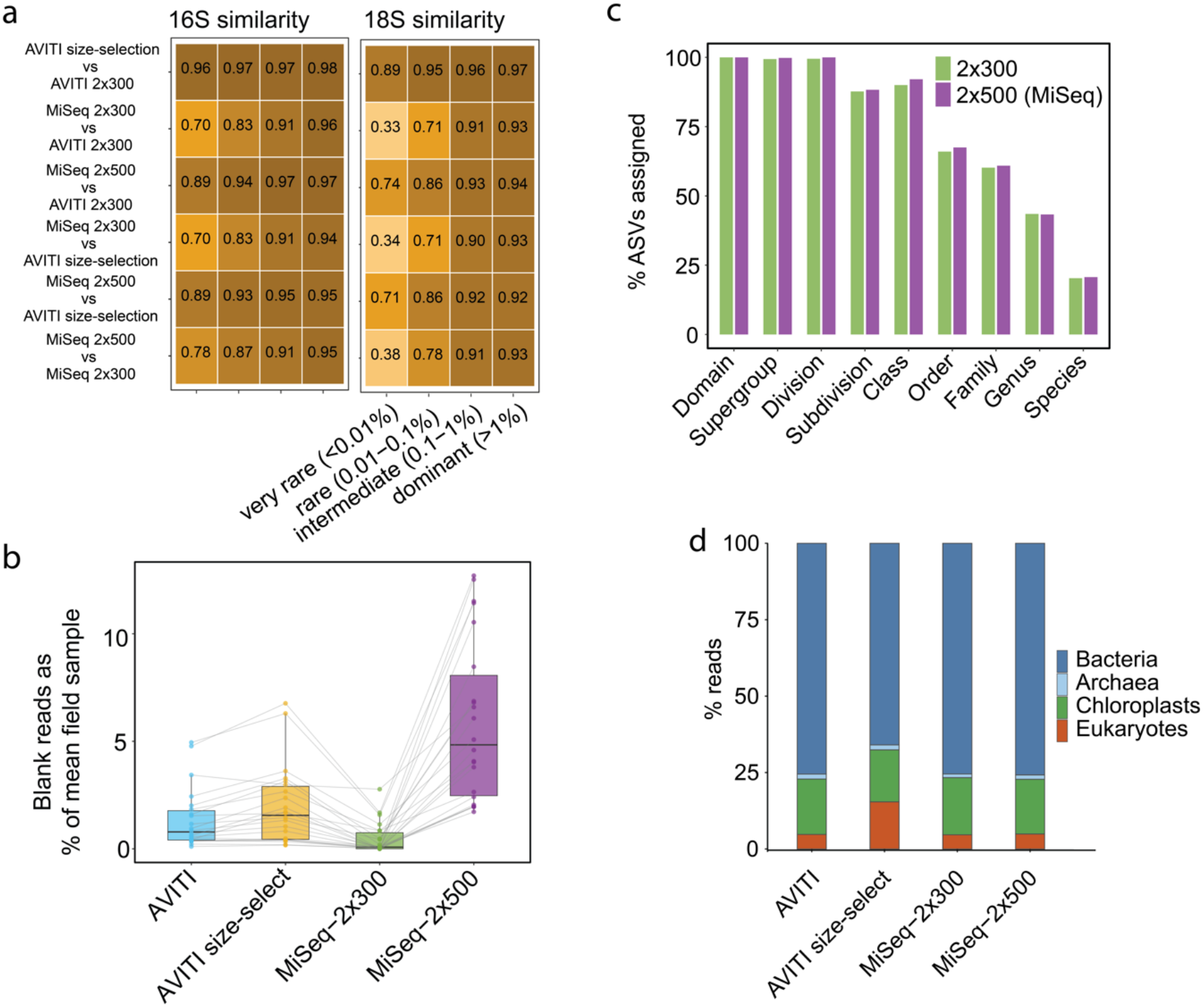
Comparison of index hopping, community data, and taxonomy assignment across platforms and chemistries. a, 16S and 18S Bray-Curtis similarity between all pairwise sequencing platform comparisons, across four abundance categories (very rare, <0.01%; rare, 0.01–0.1%; intermediate, 0.1–1%; dominant, >1%). Taxonomic classes were assigned to a single abundance category based on their mean relative abundance across all four sequencing runs (adapted from Logares et al., 2014). b, Percent of blank samples relative to mean of field samples as a proxy for index hopping across four sequencing runs. The line connecting the points connects the same blank across the different sequencing runs. c, Percentage of ASV sequences that are classified via sklearn and PR2 for the subset of matching forward and reverse fragments of 2×300 vs. the full sequences of 2×500. For this comparison, 2×300 ASVs from all sequencing runs were combined to maximize the number of comparisons, though the results were highly similar when only using MiSeq 2×300. d, Overall read proportions of the entire run of all samples across all sequencing runs.

Lastly, we considered 18S recovery and two potential means of improving it. First, we examined the outcome of the overlapping sequences of MiSeq 2×500 on 18S diversity and classification compared to the 2×300 chemistries. To compare 2×500 directly with 2×300, we restricted the 2×500 ASVs to those 3,105 ASVs that matched exactly between the stitched, non-overlapping forward–reverse 18S ASVs and the full 2×500 ASVs. 98.7% had a one-to-one match in the 2×500 data, indicating that the longer overlapping reads did not improve ASV resolution. Among the 3,105 ASVs, taxonomy assignment was increased by 0–2.1% across taxonomic levels (Fig. 2c) in the 2×500 ASVs. Thus, for the V4-V5 region of the 18S, 2×500 sequencing does not meaningfully improve taxonomy or ASV resolution, despite closing the sequencing gap left by 2×300 sequencing. Second, we evaluated the performance of the gel extraction approach. As previously mentioned, the excised 16S and 18S pools were recombined at a 50:50 molar ratio, vs 17% 18S in the original, non-size-selected pool. Upon sequencing, the overall proportion of 18S sequences in the size-selected vs non-size-selected libraries was 15.42% vs 4.77% ± 0.10% (Fig. 2d). As indicated from the small standard deviation in the non-size selected runs, it appears that the different platforms and chemistries yielded similar 18S proportions in both field samples and mock communities. Nevertheless, while the amount of 18S was still strongly biased against during sequencing, it was less biased against relative to 16S in the size-selection (3.24-fold bias, 15.42% sequenced vs 50% titrated) vs original (3.56-fold bias, 4.77% sequenced vs 17% original), suggesting that samples with higher 18S are less biased against. We observed no apparent differences in the taxonomic differences in either prokaryotes or eukaryotes between size-selected and non-size-selected approaches (R^2^=0.996 for 16S, 0.988 for 18S; Supplementary Figure 2).

## CONCLUSIONS

Across all sequencing chemistries, ecological patterns were highly conserved despite technical differences (e.g., index-hopping in MiSeq 2×500); these differences, together with price and throughput, can inform platform choice. Our approach also demonstrated that bias against 18S is conserved similarly by all of the platforms, but that with our size-selection approach, this can be adjusted for, with no detectable bias against taxa. In conclusion, while platform-specific issues remain to be addressed, our dataset of mock communities and field samples is a valuable resource and benchmark for evaluating them, and the size-selection approach offers a path to improved 18S recovery from universal primers.

## ACKNOWLEDGEMENTS

We thank the ElbeXtreme consortium, especially Eric Achterberg, Saranya Jayachandran, Björn Raupers, Ingeborg Bußmann, Götz Flöser and the crews of R/V Littorina (GEOMAR), Prandtl (HEREON), Mya II (AWI) that facilitated sample collection, and the Hamburg Port Authority. We also thank Kerstin Petersen and Jan Muschiol for support, and Osvaldina Soares for help with DNA extractions. Further support came from the Helmholtz Association, and a Helmholtz Young Investigator Grant to D.M.N. This research was supported in part through high-performance computing resources available at the Kiel University Computing Centre. This work was supported by the DFG Research Infrastructure NGS_CC (project 407495230) as part of the Next Generation Sequencing Competence Network (project 423957469). We acknowledge funding by grant no. 03F0954A of the Federal Ministry of Research, Technology and Space (BMFTR) as part of the DAM mission “mareXtreme”, project ElbeXtreme. The authors acknowledge use of AI for code suggestions and minor polishing of the manuscript text. Outputs were reviewed and edited by the authors, who take full responsibility for the content of the manuscript.

## DATA AVAILABILITY

ASV and taxonomy tables are available via Figshare: https://figshare.com/s/16c62dbcbcae711e7185. Raw sequences are available via NCBI Project Accession PRJNA1465299 (Available upon Publication)

