## Supplemental Figures for "Illumina MiSeq, MiSeq i100, and Element AVITI yield concordant ecological patterns for universal 16S and 18S rRNA gene amplicon sequencing"

**Supplementary Figures**

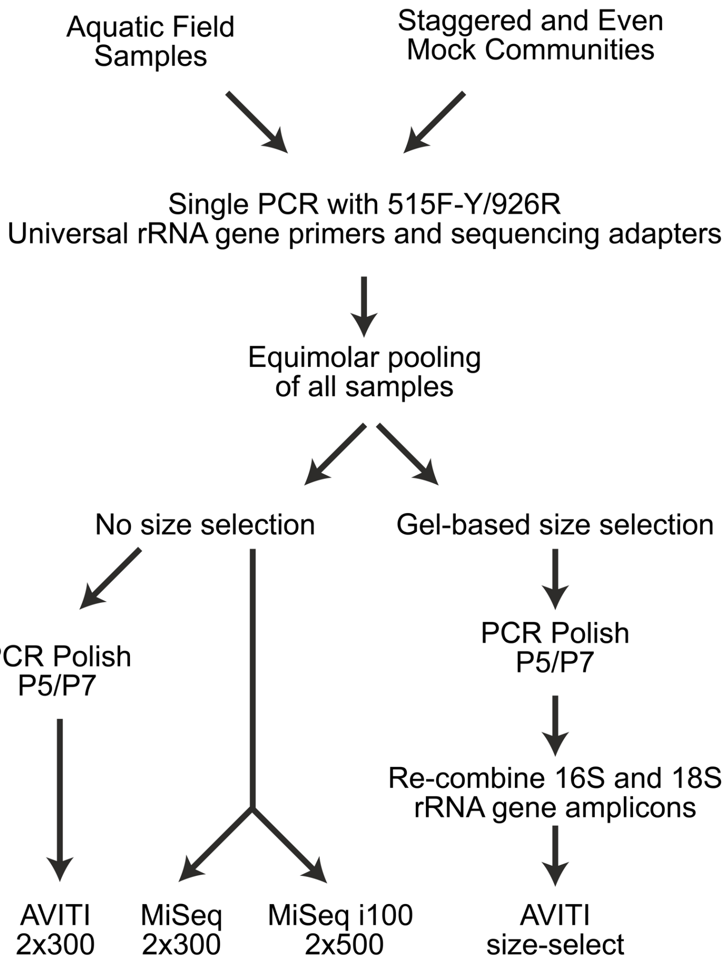


**Supplementary Figure 1.** Overview of workflow and results of size-selection and sequencing of libraries across platforms. a, Field samples and mock communities were amplified with universal primers and pooled in approximately equimolar concentrations. Then, the pooled library was split and processed with different approaches as indicated.

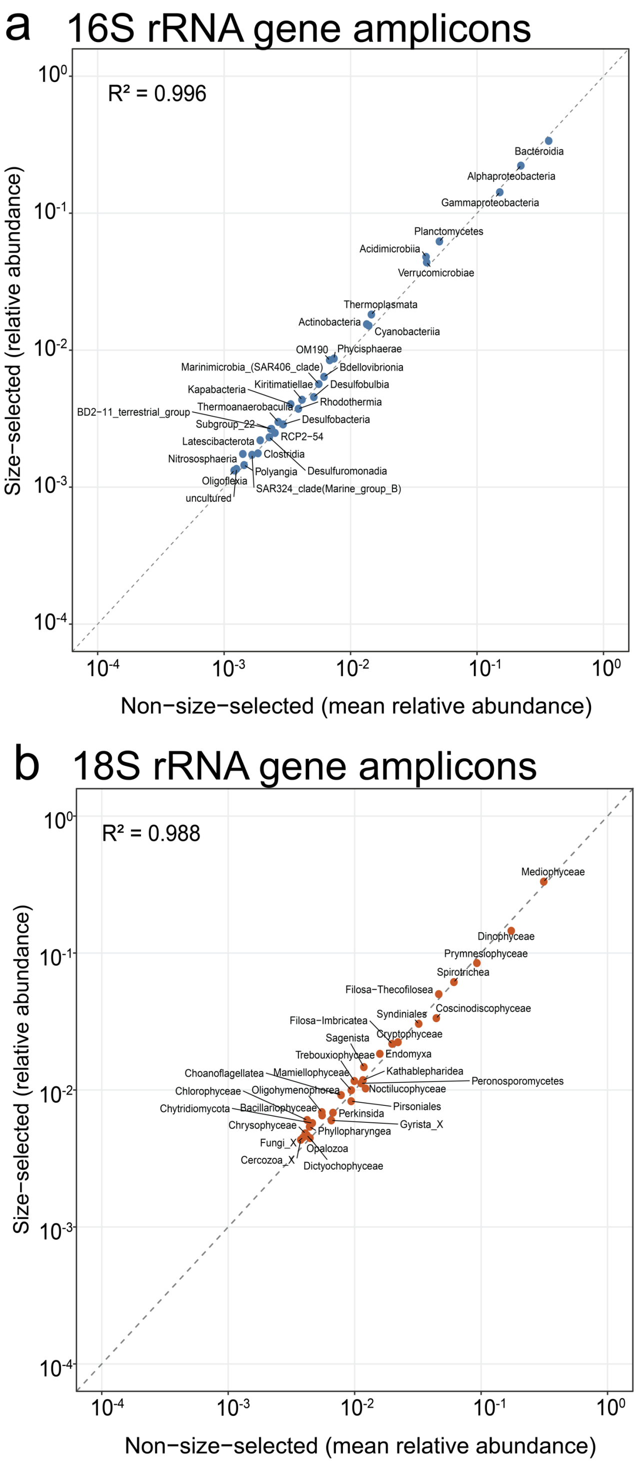


**Supplementary Figure 2.** Taxonomic composition of prokaryotic (16S, left) and eukaryotic (18S, right) communities is highly conserved between size-selected and non-size-selected approaches. Scatter plots show the top 30 Classes (relative abundance) in non-size-selected libraries (x-axis) vs the size-selected (y-axis). The dashed line indicates the 1-to-1 relationship.
